# Glucocorticoids and cortical decoding in the phobic brain

**DOI:** 10.1101/099697

**Authors:** Simon Schwab, Andrea Federspiel, Yosuke Morishima, Masahito Nakataki, Werner Strik, Roland Wiest, Markus Heinrichs, Dominique de Quervain, Leila M. Soravia

## Abstract

Glucocorticoids—stress hormones released from the adrenal cortex—reduce phobic fear in anxiety disorders and enhance psychotherapy, possibly by reducing the retrieval of fear memories and enhancing the consolidation of new corrective memories. Glucocorticoid signaling in the basolateral amygdala can influence connected fear and memory-related cortical regions, but this is not fully understood. Previous studies investigated specific pathways moderated by glucocorticoids, for example, visual-temporal pathways; however, these analyses were limited to a-priori selected regions. Here, we performed whole-brain pattern analysis to localize phobic stimulus decoding related to the fear-reducing effect of glucocorticoids. We reanalyzed functional magnetic resonance imaging (fMRI) data from a previously published study with spider-phobic patients and healthy controls. The patients received either glucocorticoids or a placebo treatment before the exposure to spider images. There was moderate evidence that patients with phobia had higher decoding of phobic content in the anterior cingulate cortex (ACC) and the left and right anterior insula compared to controls. Decoding in the ACC and the right insula showed strong evidence for correlation with experienced fear. Patients with cortisol reported a reduction of fear by 10–13%; however, there was only weak evidence for changes in neural decoding compared to placebo which was found in the precuneus, the opercular cortex, and the left cerebellum.

## 1. Introduction

Anxiety is a common disorder with a lifetime prevalence of 8%–16% in the North and South America population (Magee et al., 1996; Vicente et al., 2006). Among anxiety disorders, specific phobias are the most common, with a lifetime prevalence of 12.5%, and affect both female and male individuals of all ages (Kessler et al., 2005). Phobia can be related to fear of high altitudes, airplane travel, enclosed spaces, terror attacks, or animals such as snakes and spiders. Reactions can range from personal distress to panic attacks. To reduce the fear, most individuals with phobia avoid the stimulus or the situation, which entails considerable restrictions in their lives. Confrontation with the phobic stimulus or situation (or even its anticipation) almost invariably provokes the retrieval of past phobic memories, which consequently leads to a fear response (Cuthbert et al., 2003; de Quervain and Margraf, 2008). This mechanism supports the consolidation of additional fear memories and ultimately strengthens these fear memory traces (Sara, 2000). Therefore, the retrieval and consolidation of fearful memories seems to be an important factor for the maintenance of phobic disorders.

Evidence shows that cognitive-behavioral therapy (CBT), including exposure and cognitive restructuring, is efficacious and reorganizes processing in key regions including the amygdala, insula, and cingulate cortex (Hauner et al., 2012; Shin and Liberzon, 2010). However, up to one-third of the patients with anxiety disorders do not respond to CBT (Cuthbert, 2002; Heimberg, 2002; Hofmann and Smits, 2008). Drugs with the potential to enhance memory extinction processes are therefore promising candidates for enhancing exposure therapy. Over the last decade, various studies have demonstrated that glucocorticoids are involved in memory regulation (for an overview, see de Quervain et al., 2017). Specifically, glucocorticoids impair emotional long-term memory retrieval (de Quervain et al., 1998) while enhancing the consolidation of new memories (de Quervain et al., 2009). Previous studies demonstrated that administration of glucocorticoids reduces phobic fear in patients with anxiety disorders (Aerni et al., 2004; Soravia et al., 2006) and improves extinction-based psychotherapy (de Quervain et al., 2011; Soravia et al., 2014). Similarly, stress-induced cortisol elevation can reduce the negative affect after stress (Het et al., 2012). Thus, glucocorticoid treatment in combination with exposure therapy, is a novel and promising approach for a more effective treatment of phobia (Bentz et al., 2010).

Glucocorticoids interact with the noradrenergic system in the basolateral part of the amygdala, which has projections to temporal brain areas, such as the hippocampus (de Quervain et al., 2009). Thus, anxiolytic effects from glucocorticoids can induce changes in the amygdala that affect connected cortical areas, i.e., influence the retrieval and processing of phobic information in the temporal lobe. In our previous work, we focused on the visual temporal pathway, including the lingual gyrus, fusiform gyrus, and the amygdala (Nakataki et al., 2017). In this dynamic causal modeling (DCM) study, we found that the amygdala in phobic patients has a driving visual input mediated by the pulvinar nucleus, and that glucocorticoid administration normalizes these additional amygdala inputs. We also found that the saliency network, involving the anterior cingulate gyrus and the anterior insula, is altered in patients under glucocorticoid treatment (Soravia et al., 2018). A potential mechanism is that the anterior cingulate cortex can suppress amygdala activity during the processing of emotionally conflicting information (Etkin et al., 2006). In other words, emotional conflict is resolved through top-down inhibition of the amygdala by the cingulate cortex. In the present study, we reanalyze the same functional magnetic resonance imaging (fMRI) data as in Nakataki et al. (2017) and Soravia et al. (2018) to further investigate the relevance of the anterior cingulate cortex in processing phobic information after glucocorticoid administration in terms of neural *decoding* of phobic material, which is in contrast to activation or connectivity studies. Unlike the study by Nakataki et al. (2017) that focused only on a few regions of interest, we implemented whole-brain multivoxel pattern analysis (MVPA) and searchlight (Kriegeskorte et al., 2006), a sensitive multivariate analysis, to investigate differences in decoding of phobic material throughout the whole brain. The dataset we analyzed contained behavioral and fMRI data from a double-blind, placebo-controlled, randomized pre-registered clinical study. Spider phobic patients received glucocorticoids (20 mg of hydrocortisone) or placebo orally one hour before a picture task provoking phobic fear while fMRI images were acquired. During the experiment, participants viewed spider and non-phobic pictures and rated their experienced subjective fear.

The aim of this study was to investigate neural decoding in the phobic brain in a novel way by means of MVPA. We hypothesized that decoding of spider images in the limbic and frontal areas, including the amygdala, insula, and cingulate cortex, may be correlated with subjective fear. We also hypothesize that changes in fear may be associated with the effects of glucocorticoids in the anterior cingulate cortex. Despite these hypotheses, we decided to implement exploratory analyses to find the effects of glucocorticoids on potentially unexpected brain areas. Thus, all analyses were whole-brain analyses with appropriate control for false positives that take into account multiple testing.

## 2. Methods

### 2.1. Participants

Data were from 36 right-handed patients who fulfilled ICD-10 criteria for specific phobia for spiders and 29 healthy control participants. MRI data were taken from our previously published clinical trial, investigating the effectiveness of cortisol treatment and cognitive behavioral therapy on spider phobia treatment (Soravia et al., 2014; ClinicalTrials.gov, NCT01574014). fMRI data from this cohort has previously been published in a DCM study and a study performing independent component analysis (ICA) (Nakataki et al., 2017; Soravia et al., 2018). Male and female subjects were originally recruited via newspaper advertisements and flyers. Diagnosis of spider phobia was based on the Diagnostic and Statistical Manual of Mental Disorders, fourth edition (DSM-IV). We used a computer-based structured clinical interview (DIA-X; Wittchen and Pfister, 1997) which was based on the Composite International Diagnostic Interview (CIDI; Robins et al., 1988). We assessed fear of spiders using the German version of the Spider Phobia Questionnaire (SPQ; Watts and Sharrock, 1984) and the Fear of Spider Questionnaire (FSQ; Szymanski and O’Donohue, 1995). These questionnaires were used to assess the severity of spider phobic symptoms in the patient group and exclude healthy control subjects with elevated fear of spiders. Healthy participants were further screened by the SCL-90-R (The Symptom Checklist-90-Revised) questionnaire (Franke, 1995) to confirm that the participants had no mental illness, including spider phobia. For the healthy controls, we also confirmed no previous history of mental illness. The exclusion criteria included any of the following conditions: history of head injury, acute, or chronic medical conditions, a recent history of systemic or oral glucocorticoid therapy, psychiatric disorders other than specific phobia, psychotropic drug treatment, smoking of>15 cigarettes per day, neurological diseases, current drug or alcohol abuse, pregnancy, use of hormonal contraceptives, current behavioral therapy, and any contraindications to MRI. Female participants were evaluated during the luteal phase of their menstrual cycle as previous studies showed that amygdala activation in response to psychological stressor depends on menstrual cycles (Chung et al., 2016) and cortisol response to stress in females during the luteal phase is comparable to males (Kirschbaum et al., 1999). After diagnostic assessment, the spider-phobic patients were randomly assigned to two groups according to a double-blind, placebo-controlled design.

We carefully performed data quality checks and excluded three individuals in the cortisol group, three in the placebo group, and five controls from data analysis due to incomplete fMRI or behavioral data (12% excluded in total), or head movements larger than 4 mm of translation or 4 degrees of rotation (5% excluded); for details, see Supplementary Fig. 1A. Compared to our previous studies (Nakataki et al., 2017; Soravia et al., 2018), we excluded two more patients in the control and two more patients in the placebo group due to head movements; the control group had the same number of subjects. The reason for being stringent and excluding individuals with head movements is that MVPA is based on a first level GLM, and movement spikes can introduce spurious effects (Power et al., 2012). After exclusions, we analyzed a final sample of 54 participants: 15 patients in the cortisol group, 15 patients in the placebo group, and 24 healthy controls (see Table 1 for demographic details). After providing a complete description of the study to the participants, written informed consent was obtained. The study was approved by the ethics committee of the Canton of Bern, Switzerland (Nr. 161/07) in accordance with the principles of the Declaration of Helsinki and the Swiss authority for pharmaceutical drugs (Swissmedic) and was registered at ClinicalTrials.gov (Soravia et al., 2014; ClinicalTrials.gov, NCT01574014). All patients were offered to attend an exposure-based short-term group therapy after the study.

**Table 1.** Demographics, baseline fear and cortisol levels.

|  | Patients with<br>cortisol<br>(*n* = 15) | Patients with<br>placebo<br>(*n* = 15) | Healthy controls<br>(*n* = 24) |
| --- | --- | --- | --- |
| Median age (IQR) | 28 (24–40) | 29 (22–42) | 27 (24–29) |
| Gender (male/female) | 5/10 | 3/12 | 11/13 |
| Median BMI (IQR) | 22.2 (20.9–24.6) | 22.7 (21.3–25.0) | 21.9 (20.8–23.8) |
| Median (IQR) fear from<br>Spiders at baseline: |  |  |  |
| FSQ | 75.0 (69.5–91.5) | 77.0 (67.8–84.8) | 9.0 (2–16.5) |
| SPQ | 22.0 (20.0–26.0) | 21.0 (18.3–22.0) | 4.5 (2.8–6.0) |
| Median (IQR) cortisol concentration of<br>saliva (nmol/l): |  |  |  |
| baseline before administration | 9.0 ( 5.9–11.9) | 7.1 (6.7–12.5) | – |
| 60 min. after administration | 46.7 (17.0–56.2) | 7.6 (4.9–10.1) | – |
| 120 min. after administration | 38.6 (22.9–75.3) | 4.6 (3.0–5.8) | – |

BMI: Body Mass Index; FSQ: Fear of Spider Questionnaire; IQR: Interquartile range; SPQ: Spider Phobia Questionnaire

### 2.2. Design and procedure

Experiments were conducted at the Bern University Hospital between 2 PM and 5 PM each day. Patients and healthy controls underwent the same experimental procedure except for the diagnostic interview, substance administration, and collection of saliva samples, which only included the patients. Saliva samples were collected to control the effectiveness of the cortisol administration. Upon arrival, participants were informed about the procedure, asked to fill out the Spielberger State Anxiety Inventory (Spielberger et al., 1970), and rate their actual subjective fear, physical discomfort, and avoidance behavior on a visual analogue scale (VAS) ranging from 0 (no symptoms) to 100 (maximal symptoms). The first saliva sample was collected in the patient group using a Salivette (Sarstedt Inc., Rommelsdorf, Germany). Participants were instructed regarding the picture task, and performed a few practice trials on the computer to familiarize themselves with the rating procedure. After the oral administration of cortisol (20 mg hydrocortisone; Galepharm, Küsnacht, Switzerland) or placebo (Galepharm; lactose monohydrate, Ceolus PH 102, croscarmellose sodium, magnesium stearate) to the patients, a resting period of 30 minutes followed. To control for the cortisol level increase, sixty minutes after substance administration the second saliva sample was collected, before the beginning of the fMRI task. Functional MRI images were acquired during the picture task (24 minutes). After the scanning session, the third saliva sample was collected from the patients; therefore, the level of cortisol was measured at three time points. All participants completed questionnaires regarding anxiety (FSQ and SPQ) before the scan. Additionally, participants were asked to retrospectively rate experienced fear on a visual scale from 0 (no fear) to 100 (maximum fear) while looking at the spider images in the scanner. A further questionnaire was about side effects and about whether the patient believed that he/she had received cortisol or placebo. The saliva samples were stored at -20°C until required for biochemical analysis.

### 2.3. Paradigm

During the event-related experiment, participants viewed 80 randomized pictures from the International Affective Picture System (ISAP; Lang et al., 2008). We presented four categories (20 trials each) of images: phobic (spiders), other animals, negative, and neutral. The presentation time was five seconds, with inter-stimulus intervals (ISI) between 10.1–13.7 s (Supplementary Fig. 1B). All participants rated their subjective fear after each trial on an analogue scale between 1 (no fear) and 4 (maximum fear).

### 2.4. Statistical analysis

Baseline variables and anxiety self-ratings were analyzed using Kruskal-Wallis and Wilcoxon rank-sum tests because of the small sample size and the distribution of the data. Cortisol levels were investigated using a 2 × 3 repeated measures ANOVA (group cortisol vs. placebo, and 3 time points). MVPA was performed with the “Searchmight” toolbox (Pereira and Botvinick, 2011) and a nonparametric ANOVA with SnPM (v. 13.1.05). For correlations, Spearman’s rho correlation was used. Statistical analysis was performed in R (v. 3.4.4; R Development Core Team, 2009). All voxel-wise tests were corrected for multiple comparisons using FDR (false discovery rate) of 0.05 (Genovese et al., 2002). The analysis pipeline is illustrated in Supplementary Fig. 1C. For interpretation of *p*-values in the abstract and discussion sections, we use the terminology “strong evidence” for *p* < 0.001, “moderate evidence” for 0.001 ≤ *p* < 0.01, and “weak evidence” for 0.01 ≤ *p* < 0.05; and all tests were two-tailed.

### 2.5. Hormone analysis

We analyzed free cortisol concentrations in saliva using commercially available chemiluminescence immunoassays (cortisol: CLIA; IBL-Hamburg, Germany). The inter-and intra-assay coefficients of variation were below 10%. The samples of all subjects were analyzed in the same run to reduce error variance caused by intra-assay variability.

### 2.6. MRI data acquisition and pre-processing

Functional images were acquired with a 3T Siemens Magnetom Trio and a 12-channel head coil, using an interleaved EPI sequence (579 volumes; 37 slices; voxel, 3.6 × 3.6 × 3 mm^3^; gap thickness, 0 mm; matrix size, 64 × 64; FOV, 230 × 230 mm^2^; TR, 2500 ms; TE, 30 ms). For structural images, a high-resolution 3D T1-weighted imaging protocol (modified driven equilibrium Fourier transform, MDEFT) was used (176 sagittal slices; thickness, 1.0 mm; FOV, 256 × 256 mm^2^; matrix size, 256 × 256; voxel, 1 × 1 × 1 mm^3^; TR, 7.92 ms; TE, 2.48 ms). Pre-processing is illustrated in Supplementary Fig. 1C. We performed standard pre-processing using SPM8 (http://www.fil.ion.ucl.ac.uk/spm) and normalized to MNI space (2 × 2 × 2 mm^3^), except that data were not smoothed due to subsequent pattern analysis.

### 2.7. Multivoxel pattern analysis

We used whole-brain multivoxel pattern analysis (searchlight MVPA) with a classifier to investigate individual stimulus decoding on the subject level (Kriegeskorte et al., 2006). This step was performed within-subjects for all individuals to identify brain voxels that contain information to classify between phobic images and non-phobic images (negative, animal, and neutral images). The resulting classification accuracy maps, one per subject, represented brain areas that decoded phobic content versus the three other categories and contained values from 0–1, with 1 indicating 100% accuracy in classification of the phobic images. A major benefit of MVPA is the increased power to detect cognitive states (Haynes and Rees, 2006; Norman et al., 2006) compared to the standard mass-univariate approach. Multivariate approaches use the BOLD signal from multiple voxels (patterns) to predict stimulus category. Therefore, we can detect smaller regional differences compared to classical approaches. This shifts the interest of whether a single voxel responds differently in two conditions (mass-univariate) to whether a pattern of activity carries enough information to distinguish between the two conditions (multivariate). Prior to MVPA, a GLM was performed for each trial, including a regressor for the single trial and a regressor coding for all remaining trials, which is best practice for the subsequent MVPA (Mumford et al., 2012). We also included a CSF (cerebrospinal fluid), a WM (white matter), and six movement parameters and their first-order derivatives in the model as nuisance regressors. The resulting beta estimate maps of the individual trials were subjected to a whole-brain MVPA using a searchlight approach (Kriegeskorte et al., 2006) that involved a Gaussian Naive Bayes classifier with leave-one-sample-out cross-validation (Pereira and Botvinick, 2011). In other words, the testing set (a single image) was independent from the training images. The searchlight involved a cube of 3 × 3 × 3 voxels (6 × 6 × 6 mm^3^). Classification was performed between 20 spider pictures and 20 randomly sampled pictures from the three other categories (negative, animal, and neutral). The classification step was performed 60 times (bootstrapping) using the same 20 spider images but always a different randomly sampled set of 20 negative, animal, and neutral pictures (with a random image left out for testing) and a whole-brain mean accuracy map was created for each subject. As already stated, this accuracy map is the average percentage (across the bootstrapping) of correct classification of spider pictures versus that for the three other categories, and areas above chance accuracy can simply be interpreted as the areas for the neuronal decoding of spider pictures.

First, we performed a voxel-wise one-way ANOVA with the three group labels as levels (non-parametric permutation test with SnPM) and a FDR correction of 0.05 to identify regions that differed between the three groups; data were smoothed beforehand (FWHM = 8 mm). This is the classical analysis approach but does not take into account the high spatial resolution in MVPA. In other words, this analysis was to test the general hypothesis that fear-related regions are involved in the processing of phobic content, i.e. to validate that we actually measure decoding of phobic stimuli.

To investigate the treatment effect, we compared the accuracy maps (FWHM = 2 mm) of the placebo and cortisol group in a two-sample test using FSL “randomize” with 5,000 permutations and applied TFCE (threshold free cluster enhancement). However, it is important to note that group analyses in MVPA can be challenging as large smoothing kernels are not appropriate as these remove the spatial information. When, for example, patterns are located within a relatively large structure such as the cingulate cortex or the insula, but the patterns of activation do not spatially overlap across individuals, group significance tests will not detect alignment across individuals. Such canonical analysis approaches do not take into account these individual spatial differences; hence summed binary significance maps have been suggested (Pereira and Botvinick, 2011).

Therefore, we additionally performed this analysis for the cortisol and placebo groups. We corrected the unsmoothed individual decoding maps with a FDR (0.05) correction to get above chance decoding accuracies and applied a small smoothing kernel (FWHM = 2 mm); in these significant decoding maps, the average above chance decoding accuracies for the 30 patients was 72% (range: 66% to 85%). We binarized the individual FDR corrected accuracy maps and summed them for each patient group, resulting in two count maps demonstrating brain areas that exhibited significant individual decoding in a specific number of subjects. To compare decoding between the cortisol and the placebo groups, we subtracted the two maps, resulting in a brain map with relative increase/decrease in the number of subjects with significant decoding (difference map). To threshold this map and test for significant differences, we performed a permutation test: we randomly sampled two groups, each *n* = 15, from the pooled cortisol and placebo sample, with each group containing randomly drawn individuals from both groups. We summed the maps and created two count maps, one for each group, and subtracted the maps to create a difference map. We performed this k = 500 times to create a distribution under the null hypothesis of no difference between the cortisol and placebo group. To control the type I error, we determined the upper and lower percentiles (2.5% and 97.5%); thus, only in 5% of the cases an observed result would be equal to or more extreme under the null hypothesis. These boundaries were determined voxel-wise and used to threshold the original difference maps. The average upper and lower alpha level thresholds across voxels were +3.6 subjects (SD: 0.51) and -3.4 subjects (SD: 0.54). We used the Harvard-Oxford cortical atlas to identify and label the significant brain regions.

### 2.8. Data availability

Preprocessed imaging data (cortical decoding maps), subjective fear, and related data to generate all the main results in the tables and figures, including a reproducible R notebook, is available at OSF https://osf.io/yr823/.

## 3. Results

### 3.1. Demographics, baseline fear and endocrine measures

Demographics are shown in Table 1. The three groups did not significantly differ in age (Kruskal-Wallis test, χ^2^ = 0.80, df = 2, *p =* .67). There were higher proportions of females in the two patient groups; however, the three groups did not differ significantly regarding gender (χ^2^ = 2.73, df = 2, *p =* .26). There was also no difference regarding body mass index (Kruskal-Wallis test, χ^2^= 1.03, df = 2, *p =* .60). The cortisol and placebo patient groups did not differ regarding spider phobia symptoms assessed by FSQ and SPQ at baseline before the experiment (Wilcoxon rank-sum test; FSQ: W = 99, *p =* .98; SPQ: W = 121.5, *p =* .29). At baseline, patients had significantly higher scores in spider phobic symptoms compared to healthy controls, who had no fear (Kruskal-Wallis test; FSQ: χ ^2^= 36.6, df = 2, *p <* .0001; SPQ: χ ^2^= 37.0, df = 2, *p <* .0001; Table 1). The cortisol group had 5.2 times higher cortisol levels at the beginning of the fMRI experiment, and 4.3 times higher levels at the end of the experiment compared to their baseline, while the placebo group showed no increase over time (Table 1), which was confirmed by a significant interaction group × time (repeated measures ANOVA; *F*_4,98_ =46.6, *p <* .0001).

### 3.2. Subjective fear

During the fMRI task, patients (cortisol and placebo groups) exhibited 2.1 times higher subjective fear in response to spider pictures compared to controls (Kruskal-Wallis rank sum test, χ^2^= 28.8, df = 2, *p <* .0001) (Fig. 1A). Patients with cortisol treatment had a 9.8% decrease in subjective fear while looking at spider pictures compared to patients with placebo (Wilcoxon rank sum test, W = 47.5, *p =* .021) (Fig. 1A). We found a 2.4 times higher subjective fear in patients with spider-phobia compared to controls when subjects were asked to retrospectively rate their perceived fear in the scanner after the experiment (χ^2^= 23.2, df = 2, *p <* .0001) (Fig. 1B), and patients with cortisol had a subjective fear reduction of 13.3% compared to the placebo group (W = 42.5, *p =* .011). The three groups did not differ with respect to subjective fear while looking at the negative pictures (χ ^2^= 4.57, df = 2, *p =* .10) (Fig. 1C).

**Fig. 1.**
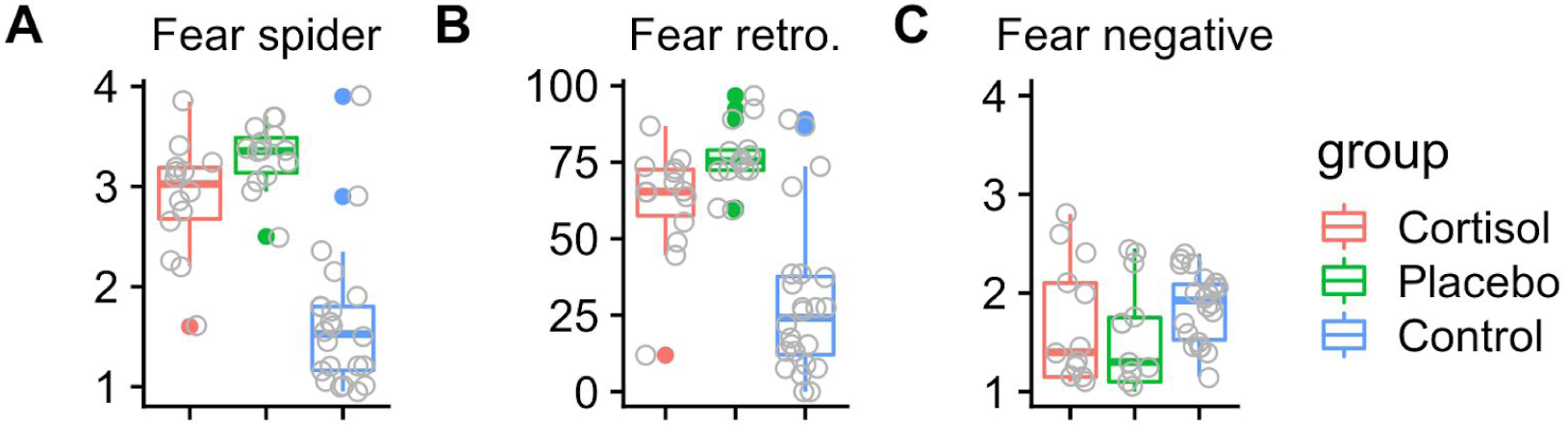
Subjective fear during the fMRI experiment. **(A)** The cortisol group exhibits a 9.8% reduction of fear towards spider pictures compared to placebo (*p =* .021). Patients (cortisol and placebo groups) had a 2.1 times higher subjective fear compared to controls (*p <* .0001). **(B)** Retrospective fear during the experiment assessed after the experiment: the cortisol group shows a 13% reduction of fear from spiders (*p =* .011). Patients had a 2.4 times higher fear compared to the controls (*p <* .0001). **(C)** No group differences were found among the three groups regarding emotional negative pictures (*p =* .10).

### 3.3. Cortical decoding of spider images

We first performed a voxel-wise test for group differences with the null hypothesis that decoding of phobic images is equal in all three groups (non-parametric one-way ANOVA, FDR 0.05). We found the strongest group effects in the left anterior insula, the right anterior insula, and in the anterior cingulate gyrus (ACC) (Fig. 2). These areas showed a higher median decoding in both patient groups (cortisol: insula left, 57%; insula right, 56%; ACC, 59%; placebo: insula left, 56%; insula right, 55%; ACC, 56%) compared to healthy controls (insula left, 50%; insula right, 50%; ACC, 50%) (Fig. 2A). It is important to note that these decoding values are averages across the areas of the anterior insula and the ACC and also across individuals; however, these decoding values are found to be much higher locally and individually, i.e., above 80% accuracy (Supplementary Fig. 2). Decoding in all three areas correlated with subjective fear rated during the scanner and explained 36%–40% of the variance (Spearman’s rank correlation; insula l., ρ = .60, *p <* .0001; insula r., ρ = .64, *p =* .0001; ACC, ρ = .64, *p <* .0001) (Fig. 3).

**Fig. 2.**
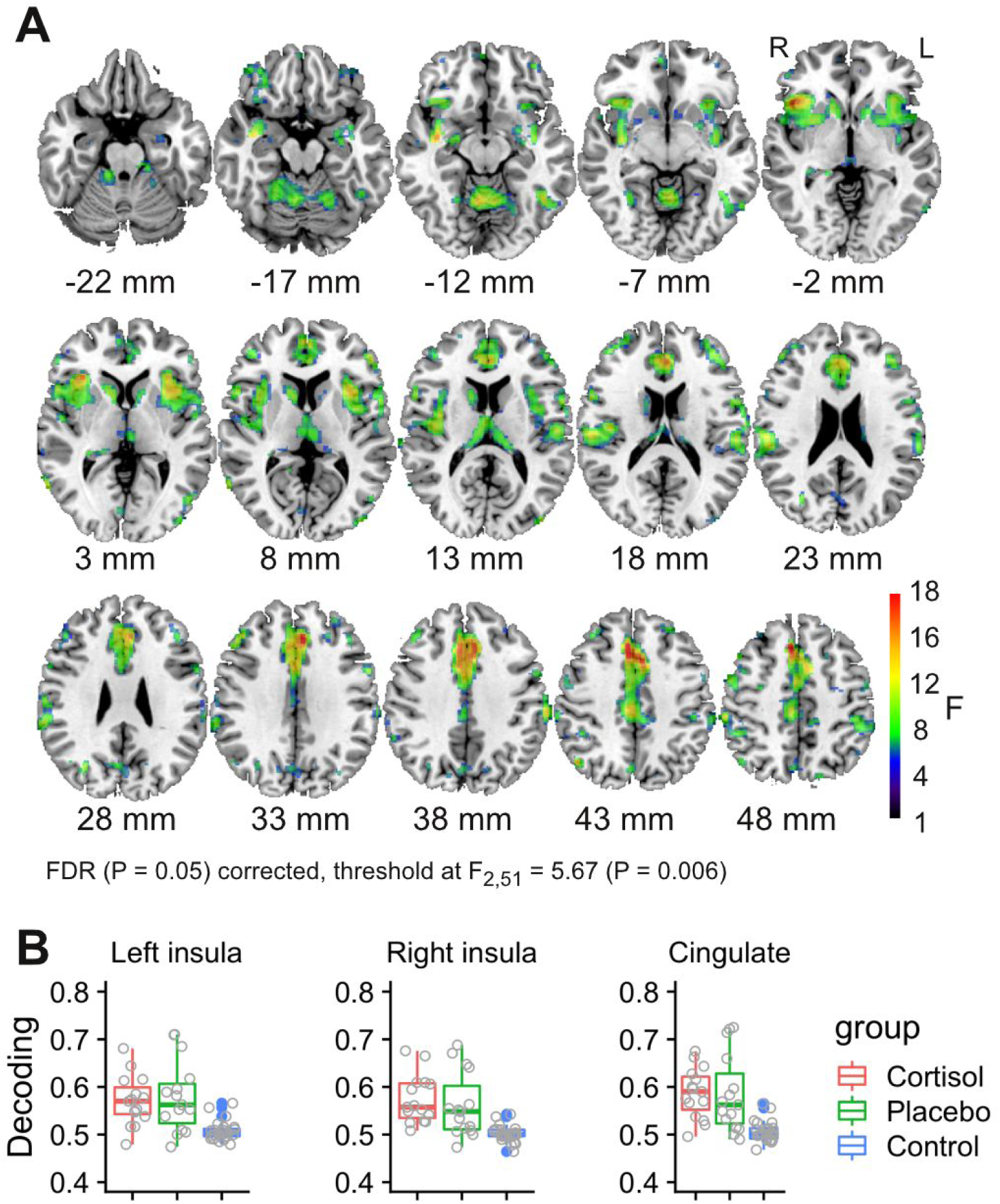
**(A)** Areas showing group effects (differences of means between the cortisol, placebo and control group using one-way ANOVA) in phobic decoding (spider vs. negative, animal and neutral images). Most prominent regions are the left anterior insula, the right anterior insula, the anterior cingulate cortex (ACC). **(B)** Post-hoc analysis showing that the group effect is due to patients (cortisol and placebo) with higher decoding accuracies compared to controls in the left anterior insula, the right anterior insula, and the ACC.

**Fig. 3.**
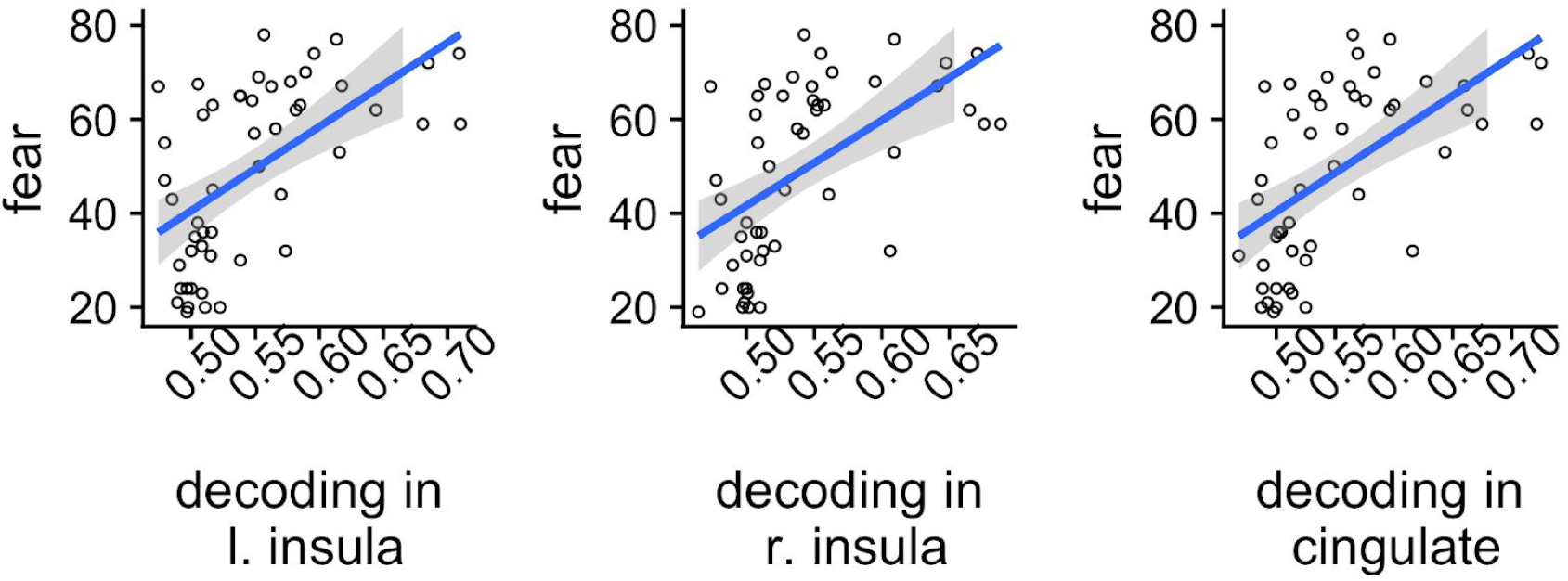
Decoding of spider images in these three regions correlated positively with subjective fear from spiders and explained 36%–40% of the variance (*R*^2^; gray areas around the regression line is the 95% confidence interval).

We compared directly by voxel-wise analysis the accuracy maps of the cortisol and the placebo group and found no significant difference for a treatment effect with respect to decoding of phobic images after applying correction for mass-univariate testing (two-sample t-test with TFCE). We performed an additional analysis to explore further significant decoding. We used the thresholded accuracy map on the subjects level (FDR 0.05 corrected) and transformed these into binary maps that we summed across individuals from the placebo and patient groups. The maps of the two groups showed high similarity (Fig. 4). We calculated a difference map with a permutation test and an alpha level of *p* = 0.05 (Fig. 5AB) (for more axial slices, see Supplementary Fig. 3). In the cortisol group, an increased number of individuals showed decoding of spider images in the right precuneus cortex (+8 individuals; 12 cortisol; 4 placebo), the left central opercular cortex (+6; 8 cortisol; 2 placebo), the left cerebellum (+7; 8 cortisol; 1 placebo), the right parietal opercular cortex (+7; 9 cortisol; 2 placebo) (see Table 2 for a complete list). In the placebo group, an increased number of individuals demonstrated higher decoding in the insula (+6; 10 placebo; 4 cortisol), the ACC (+5; 7 placebo; 2 cortisol), and the frontal pole (+6; 6 placebo; 0 cortisol) (see Table 3 for a complete list).

**Table 2.**
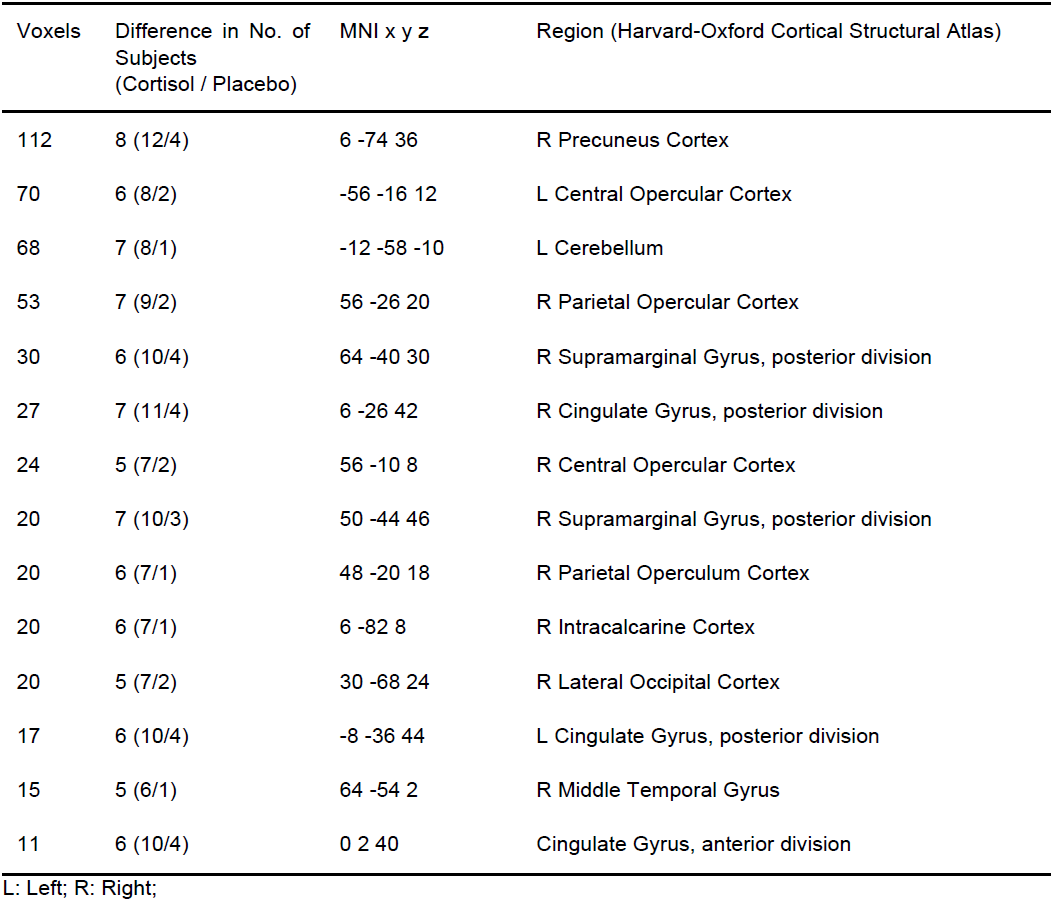
Brain regions with a significant larger number of participants in the cortisol group who have above-average decoding compared to the placebo group.

| Voxels | Difference in No. of Subjects<br>(Cortisol / Placebo) | MNI x y z | Region (Harvard-Oxford Cortical Structural Atlas) |
| --- | --- | --- | --- |
| 112 | 8 (12/4) | 6 -74 36 | R Precuneus Cortex |
| 70 | 6 (8/2) | -56 -16 12 | L Central Opercular Cortex |
| 68 | 7 (8/1) | -12 -58 -10 | L Cerebellum |
| 53 | 7 (9/2) | 56 -26 20 | R Parietal Opercular Cortex |
| 30 | 6 (10/4) | 64 -40 30 | R Supramarginal Gyrus, posterior division |
| 27 | 7 (11/4) | 6 -26 42 | R Cingulate Gyrus, posterior division |
| 24 | 5 (7/2) | 56 -10 8 | R Central Opercular Cortex |
| 20 | 7 (10/3) | 50 -44 46 | R Supramarginal Gyrus, posterior division |
| 20 | 6 (7/1) | 48 -20 18 | R Parietal Operculum Cortex |
| 20 | 6 (7/1) | 6 -82 8 | R Intracalcarine Cortex |
| 20 | 5 (7/2) | 30 -68 24 | R Lateral Occipital Cortex |
| 17 | 6 (10/4) | -8 -36 44 | L Cingulate Gyrus, posterior division |
| 15 | 5 (6/1) | 64 -54 2 | R Middle Temporal Gyrus |
| 11 | 6 (10/4) | 0 2 40 | Cingulate Gyrus, anterior division |

L: Left; R: Right;

**Table 3.**
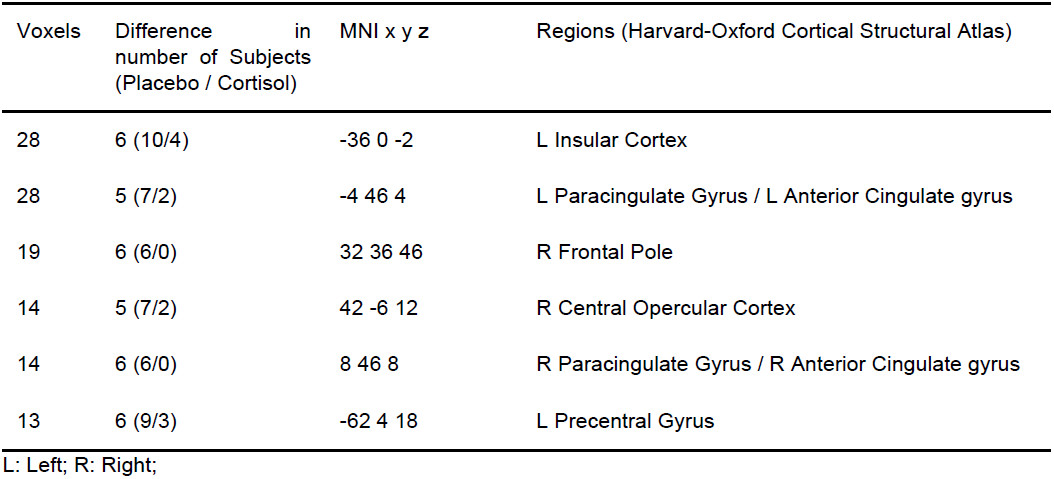
Brain regions with a significant larger number of participants in the placebo group who have above-average decoding compared to the cortisol group.

**Fig. 4.**
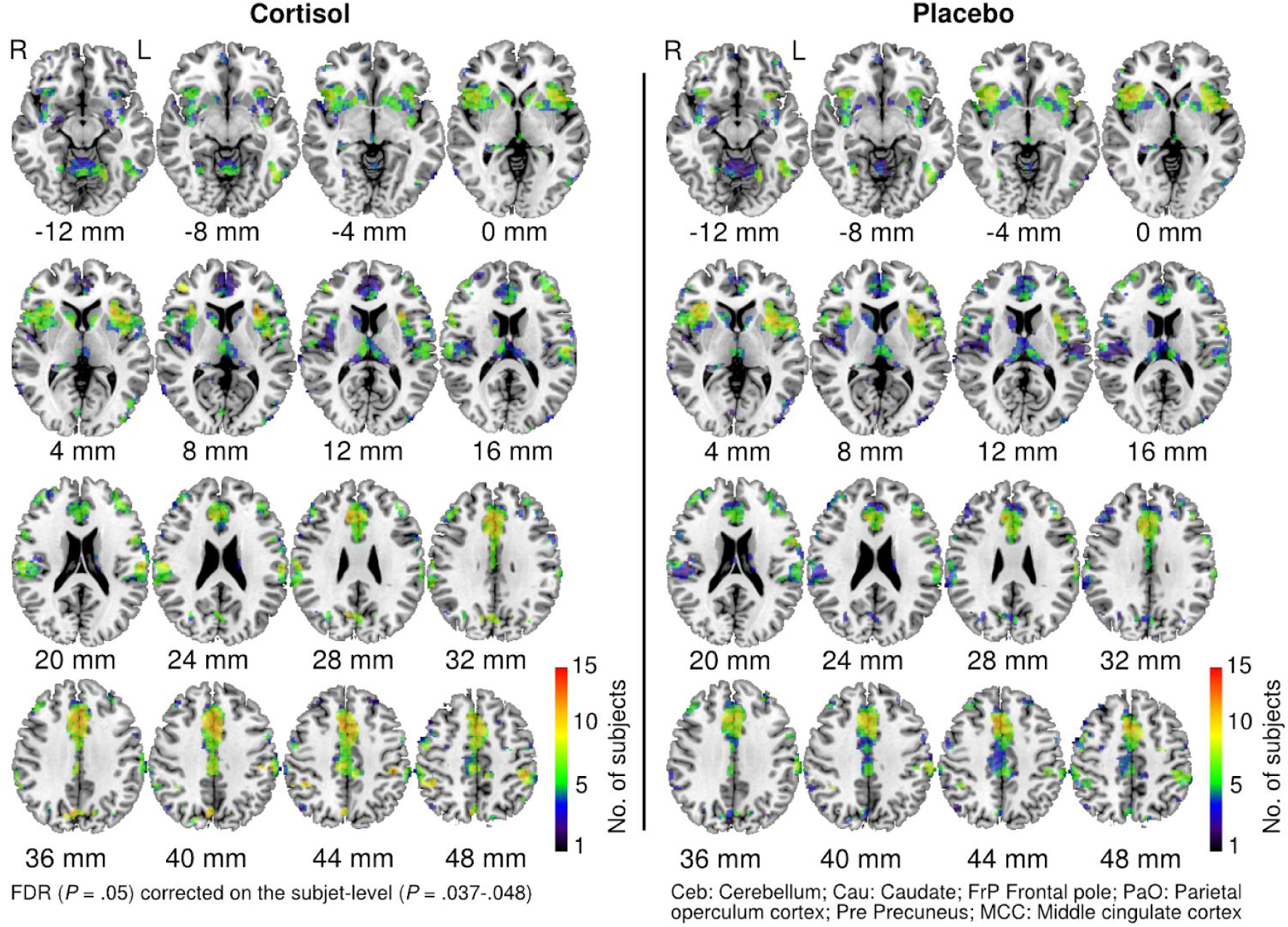
Count maps of individuals with significant decoding in the cortisol and placebo groups. Aggregated individual decoding maps (each FDR 0.05 corrected on the subject-level). Colors represent the number of subjects with significant decoding.

**Fig. 5.**
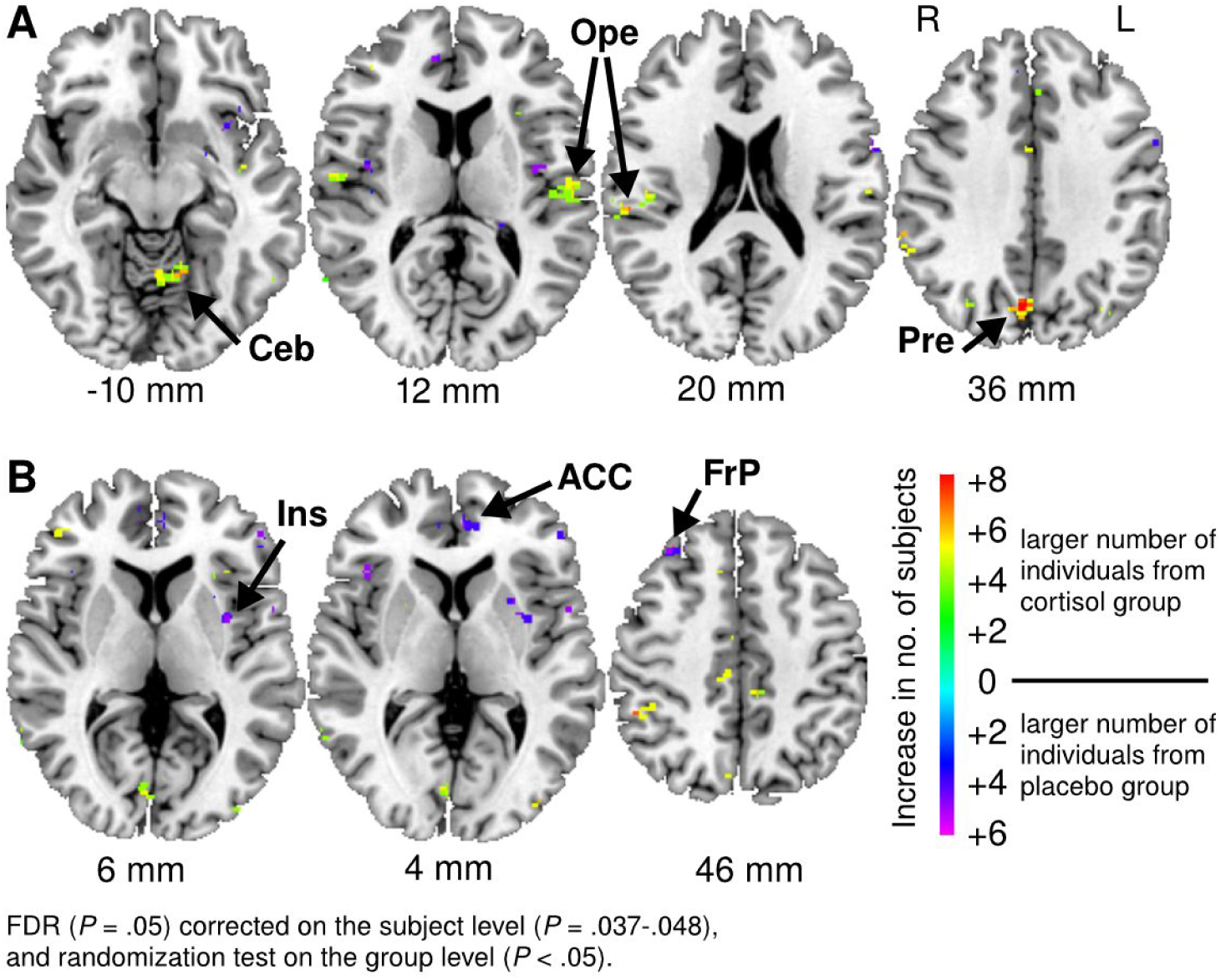
Decoding of spider images. Brain areas with a relative increase in the number of subjects who have significant individual decoding of spider images (FDR 0.05 corrected on the subject-level; randomization test *p* = 0.05 on the group level). **(A)** A larger number of individuals in the cortisol group showed decoding of spider images most prominently in the right precuneus cortex (Pre), the right and left opercular cortex (Ope), and the left cerebellum (Ceb). **(B)** A larger number of individuals in the placebo group showed decoding most prominently in the left insula (Ins), the anterior cingulate gyrus (ACC), and the frontal pole (FrP).

## 4 Discussion

The aim of this exploratory study was to perform a whole-brain multivariate pattern analysis to investigate the relevance of the anterior cingulate gyrus and other potentially relevant areas that demonstrate changes in phobic information decoding after glucocorticoid administration. Decoding of phobic images in the anterior insula and the ACC was higher in patients compared to controls (moderate evidence) and was correlated with subjective fear (strong evidence). This involvement of the limbic system and the fact that the decoding magnitudes correlated with subjective fear supports the idea that our brain-derived phenotype, the decoding (classification accuracies from MVPA), is a valid measure and closely linked to phobic information processing. These areas are well-known key players in the processing of emotions and part of the fear network (Greco and Liberzon, 2016), particularly in specific phobia (Del Casale et al., 2012), and it was shown that exposure therapy reduces activity specifically in these regions, the cingulate cortex and the insula (Hauner et al., 2012).

We compared the treatment (cortisol) and the placebo group. In general, the decoding maps of the two groups were highly similar, and the effect related to cortisol treatment seemed to have weak evidence. We performed two alternative analyses to investigate the neural signatures of the treatment effect, but these provided inconclusive results. A standard approach used for activation studies did not show any significant results. We additionally binarized the maps which was more effective for depicting patterns of individual decoding across individuals. In the placebo group with higher experienced fear, a large number of individuals showed decoding in frontal regions (weak evidence): the insula, the ACC and the right lateral frontal pole. It has been demonstrated that these regions are involved in fear and memory processes: the insula and ACC are key player in the fear circuitry and their role has been well demonstrated in anxiety disorders (Shin and Liberzon, 2010). The frontal pole is involved in various cognitive functions, such as mentalizing, multitasking, and episodic memory (Christoff and Gabrieli, 2000; Gilbert et al., 2006). It has been shown that the lateral frontal pole co-activates with the ACC and the anterior insula (Gilbert et al., 2010). This result, consistent with the symptom reduction achieved through CBT, was associated with lower responsiveness in the bilateral insula and the ACC and a reduction in cerebral blood flow (Hauner et al., 2012; Soravia et al., 2016a). Contrary, the cortisol group showed decoding in the precuneus, the anterior cerebellum, and the opercular cortex (weak evidence). The precuneus is a hub involved in multimodal, attentional and memory processes; it is also involved in the default mode network (Soravia et al., 2016b). Functional connectivity mapping of this hub showed negative connectivity with the amygdala (Zhang and Li, 2012). Thus, it may be possible that glucocorticoid related changes in the amygdala may moderate the precuneus cortex; however, this needs to be further investigated. A recent study with healthy participants showed that cortisol disrupts ventromedial prefrontal cortex functioning and its communication with other brain regions such as the cingulate cortex and parahippocampal gyrus (Kinner et al., 2016). This is a potential mechanism behind our finding that the cortisol group showed reduced decoding in the ACC. The insula is particularly involved in processing of emotionally salient information as it is strongly connected with limbic structures, the cingulate cortex, the amygdala, and prefrontal regions. Hence, the acute administration of glucocorticoids and similarly CBT may reduce the hyperactivation in response to phobic stimuli in brain regions that are crucially involved in identifying fearful stimuli and generating fear response, such as the insula (Duval et al., 2015).

In this study, we used pattern analysis, a method applied on the individual subject level. This yields individual decoding patterns that only partially overlap due to local functional decoding differences among individuals. Therefore, it can be challenging to find consistent results in standard voxel-wise group analyses, and such group analyses are generally not recommended in combination with MVPA (Stelzer et al., 2013). We addressed this issue using count maps (Pereira and Botvinick, 2011) that were thresholded on the individual, and on the group level with a randomization test, providing robust statistics but also accounting more thoroughly for individual variation in decoding.

Some limitations of this study merit discussion. First, we present data from a sample with small subgroups; however, this is commonly encountered in neuroimaging studies involving clinical samples and drug administration. We excluded some subjects due to head movements; it is well known that head movements can confound results in neuroimaging (Deen and Pelphrey, 2012; Power et al., 2012). Correlations and functional connectivity seem especially sensitive to movement and can exhibit false effects in response. While there is agreement on how to analyze standard activation studies, there are not many guideline recommendations on MVPA group analyses (Pereira and Botvinick, 2011). We addressed this problem using two alternative analyses: first, a standard voxel-wise analysis with the accuracy maps, and a second analysis of the binarized accuracy maps. A further limitation is the gender imbalance: there were 46% males in the control group, but only 33% in the cortisol and 20% in the placebo group.

To summarize, this study elucidates cortical decoding patterns of phobic stimuli related to glucocorticoid treatment. We found moderate evidence of neural decoding of phobic material in the anterior cingulate cortex and bilateral anterior insula. Concerning the acute effect of glucocorticoid administration on processing of phobic fear, there was weak evidence indicating for posterior decoding (precuneus, cerebellum, and operculum) in the treatment group (cortisol), and a frontal decoding (insula, ACC and frontal pole) in the placebo group. Taken together, the findings provide some new insight into the neuronal decoding of the phobic brain.

## Acknowledgments

This work was supported by the Swiss National Science Foundation (No. 124947, No. 171598) and by the Medical Faculty of the University of Bern (520.10). We thank Melanie Fisler, Joëlle Witmer, Basil Preisig, and Yvonne Renevey for their assistance. We thank all participants for their time and efforts.

