## Supplementary file for "Glucocorticoids and cortical decoding in the phobic brain"

### Supplementary Material

#### Supplementary Figures

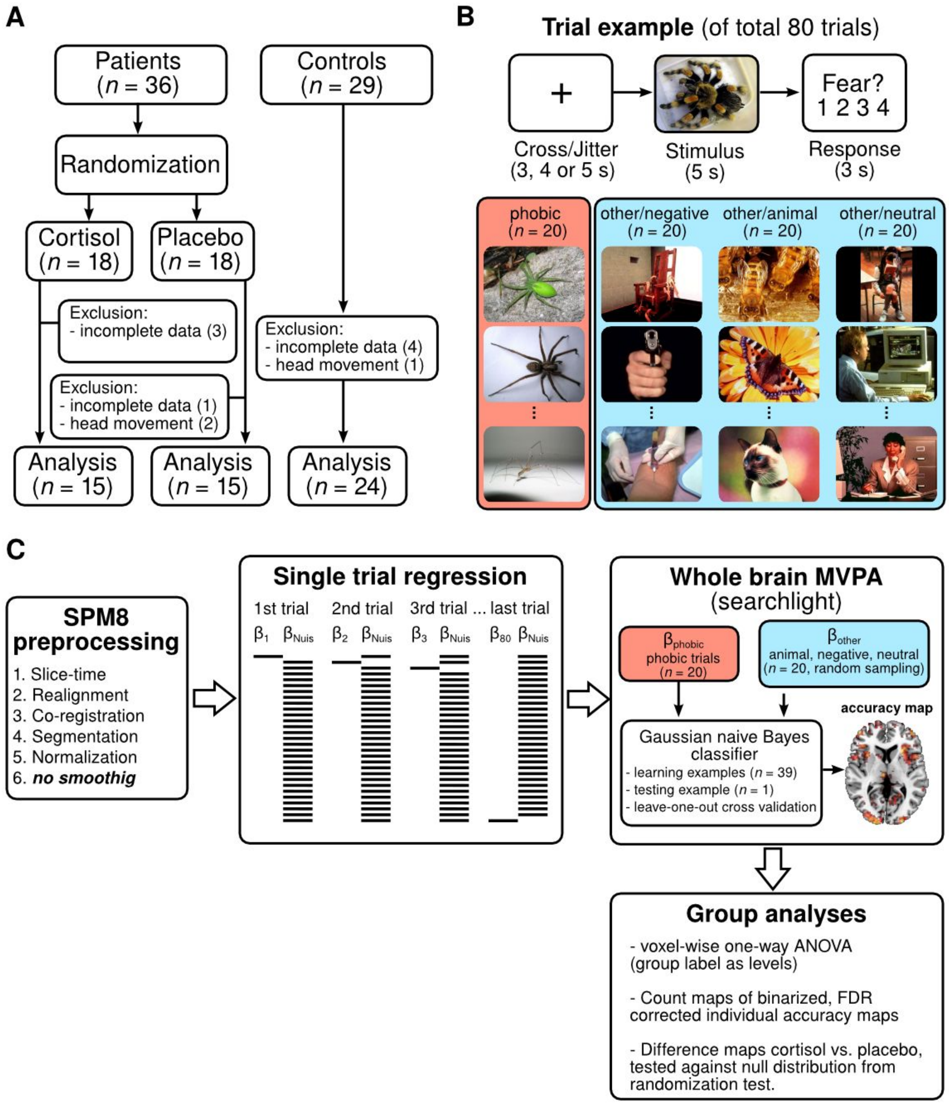


**Supplementary Figure 1**. Design, stimulus material and analysis pipeline. **(A)** We excluded participants with large head movements or who had incomplete fMRI or behavioural data, resulting in a total of 54 participants that were analysed. **(B)** Eighty trials were presented in four picture categories (20 each). To investigate phobic specific brain areas, we also presented emotionally negative content, non-phobic animals, and neutral images. **(C)** Image processing and analysis pipeline. After standard pre-processing (without smoothing), trial-by-trial estimates were used as examples for the machine learning to classify phobic vs. the other image categories (negative, neutral, and animal) resulting one accuracy map per subject for decoding of phobic material which were subjected to group analyses.

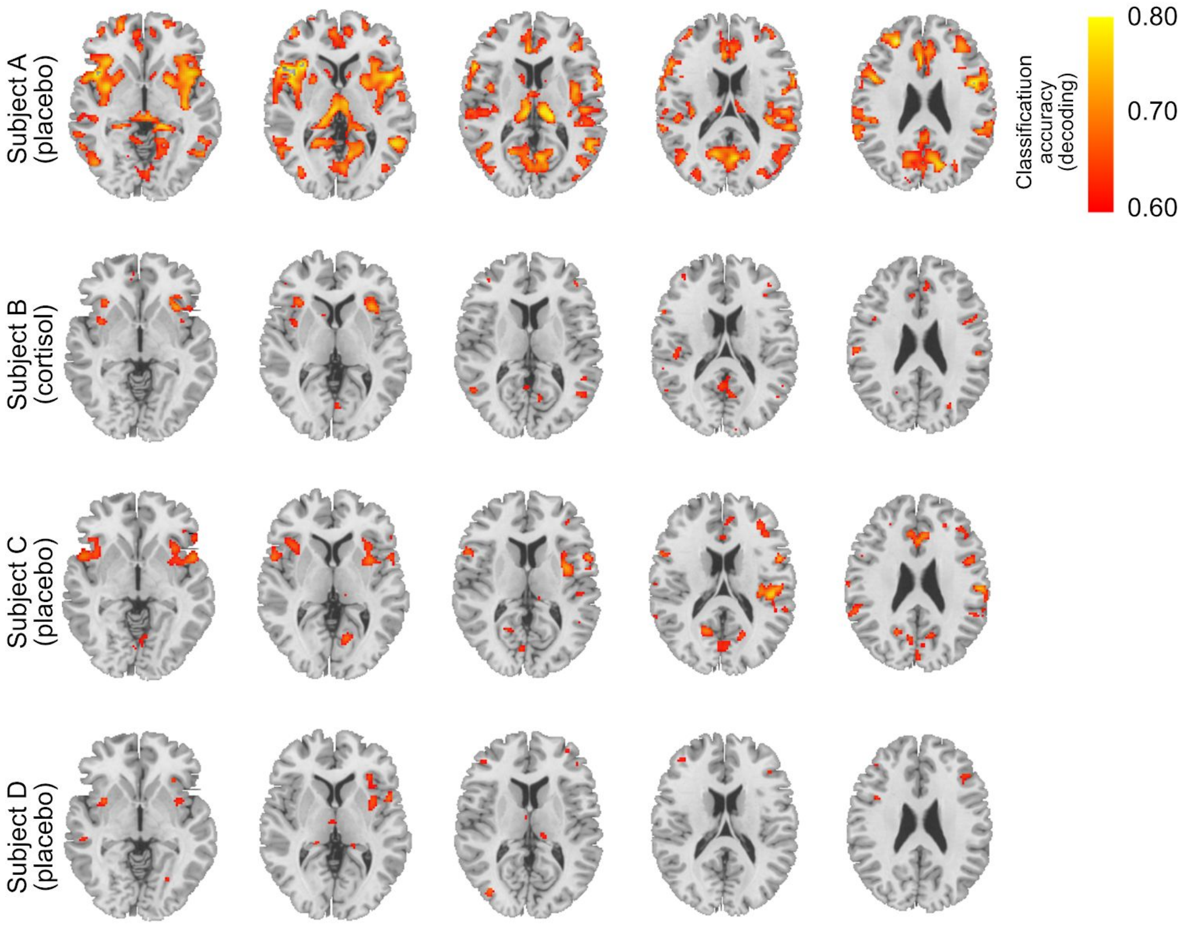


**Supplementary Figure 2**. Four random, representative individuals from the patient group (cortisol and placebo) showing individual decoding maps (classification accuracy, FDR 0.05 corrected) that can be as high as 80% in some locations of Subject A, B and C. Subject D has smaller magnitudes.

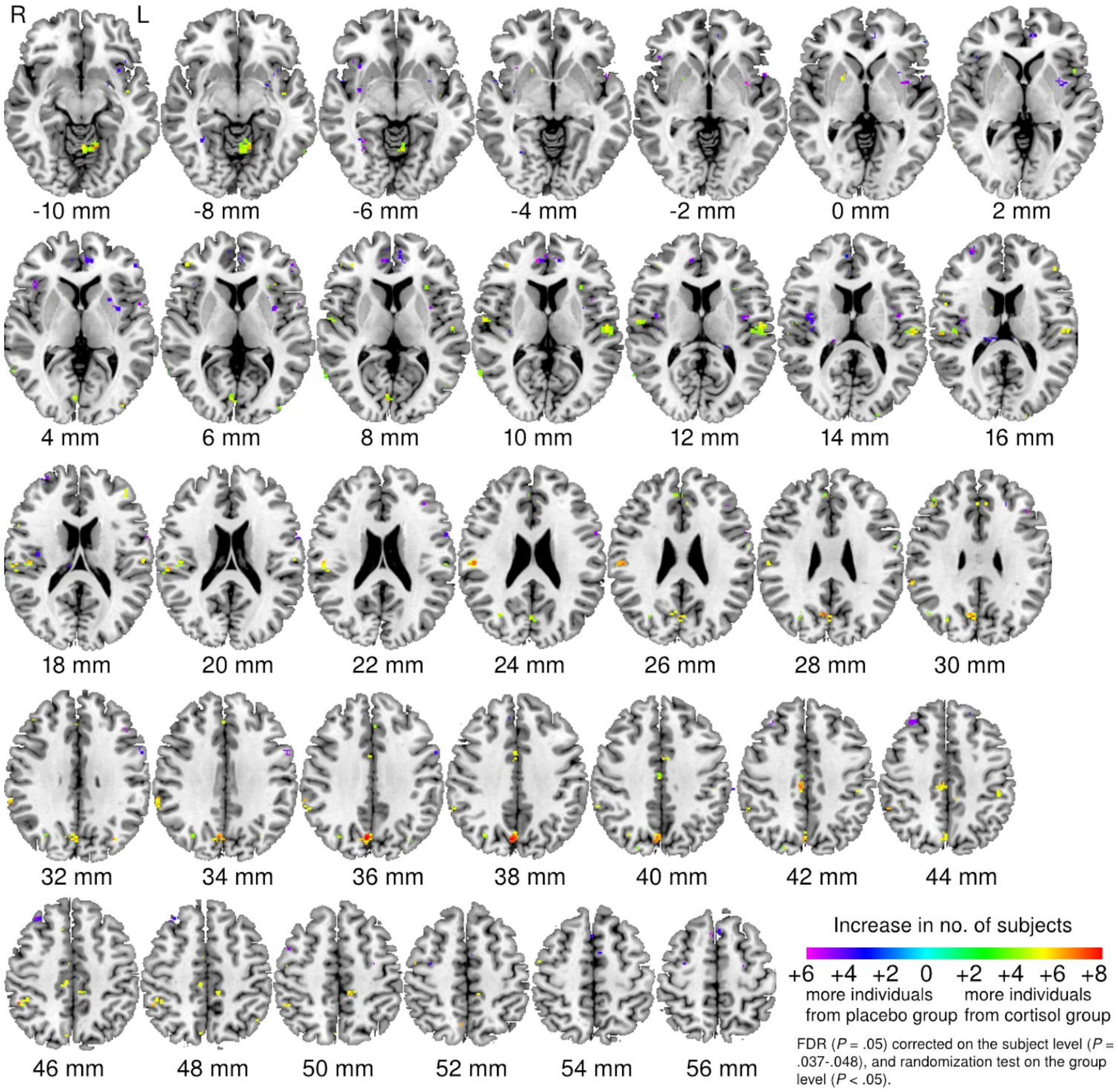


**Supplementary Figure 3**. Difference maps of decoding of spider images. Brain areas with a relative increase in the number of subjects who have significant individual decoding of spider images (FDR 0.05 corrected on the subject-level; randomization test on the group level with $p < 0.05$).
